# Cellular traits regulate fluorescence-based bio-optical phenotypes of coral photosymbionts living *in-hospite*

**DOI:** 10.1101/2023.07.06.547959

**Authors:** Audrey McQuagge, K. Blue Pahl, Sophie Wong, Todd Melman, Laura Linn, Sean Lowry, Kenneth D. Hoadley

**Affiliations:** Department of Biology, University of Alabama, Tuscaloosa, AL, United States; Dauphin Island Sea Lab, Dauphin Island, AL, United States; University of Virginia, Charlottesville, VA, United States; Reef Systems Coral Farm, New Albany, OH, United States 15

**Author notes:** Corresponding Authors: Audrey McQuagge, Kenneth Hoadley, 21.

**Keywords:** *Symbiodiniaceae*, Coral photosymbionts, Functional Traits, Chlorophyll-*a* fluorescence, Phenomics

## Abstract

Diversity across algal family *Symbiodiniaceae* contributes to the environmental resilience of certain coral species. Chlorophyll-*a* fluorescence measurements are frequently used to determine symbiont health and resilience, but more work is needed to refine these tools and establish how they relate to underlying cellular traits. We examined trait diversity in symbionts from the generas *Cladocopium* and *Durusdinium,* collected from 12 aquacultured coral species. Photophysiological metrics (Φ_PSII_, σ_PSII_, ρ, τ_1_, τ_2_, ABQ, NPQ, and qP) were assessed using a prototype multi-spectral fluorometer over a variable light protocol which yielded a total of 1360 individual metrics. Photophysiological metrics were then used to establish four unique phenotypic variants. Corals harboring C15 were predominantly found within a single phenotype which clustered separately from all other coral fragments. The majority of *Durusdinium* dominated colonies also formed a separate phenotype which it shared with a few C1 dominated corals. C15 and D1 symbionts appear to differ in which mechanisms they employ to dissipate excess light energy. Spectrally dependent variability is also observed across phenotypes that may relate to differences in photopigment utilization. Cell physiology (atomic C:N:P, cell size, chlorophyll-*a*, neutral lipid content) was also assessed within each sample and differ across phenotypes, linking photophysiological metrics with underlying primary cellular traits. Strong correlations between first– and second-order traits, such as Quantum Yield and cellular N:P content, or light dissipation pathways (qP and NPQ) and C:P underline differences across symbiont types and may also provide a means for using fluorescence-based metrics as biomarkers for certain primary-cellular traits.

## Introduction

The dinoflagellate algal family *Symbiodiniaceae* is genetically and phenotypically diverse, having evolved to occupy numerous niches and lifestyles, which include free-living open ocean as well as endosymbiotic roles most notably in stony corals (Muscatine & Hand, 1958; Loeblich & Sherley, 1979; Takabayashi et al., 2012). Within the coral holobiont, *Symbiodiniaceae* live within coral gastrodermal cells, in which they recycle coral waste products and in turn produce up to 90% of the coral’s energy stores through fixed carbon (Yellowlees et al., 2008). Their endosymbiotic relationship with stony corals is complex and dynamic, with differing life histories, coral hosts, and environments giving rise to vast diversity in survival strategies and physiology across the family (Wiedenmann et al., 2012; Pasaribu et al., 2016; Suggett et al., 2017; LaJeunesse et al., 2018). Elevated seawater temperatures above a certain threshold have increased the risk of coral endosymbiont loss (coral bleaching) and its associated sublethal and lethal effects worldwide (Hoegh-Guldberg & Smith, 1989; Steen & Muscatine, 1987; Tchernov et al., 2011). Coral bleaching mitigation and reef restoration relies on improved holistic understanding of the coral holobiont (the coral host and its microbiome, including endosymbiotic algae), to determine what traits underpin resilience under environmental stressors. Association with adaptable and resilient symbiont species is thought to be an important predictor of coral resilience (Suggett et al., 2017), but the underlying phenotypic differences across the family are not fully resolved.

Inherent functional traits of both the algal symbiont and the coral host, as well as their interactive physiology, govern coral bleaching susceptibility which varies across different host-symbiont combinations (Sampayo et al., 2008; Krueger & Gates 2012; Scheufen et al., 2017). Certain genera of *Symbiodiniaceae*, such as *Durusdinium* (Clade D) are often linked with higher thermal tolerance across many Caribbean and Pacific coral species where mixed assemblages of photosymbionts are common (Baker et al., 2004; Fabricius et al., 2004; LaJeunesse et al., 2003; Sampayo et al., 2008). For these ‘symbiont generalist’ coral species, dominance by the species *Durusdinium trenchii* is particularly notable among colonies that display higher thermal tolerance. Other coral species only associate with a single symbiont species (specialists), and their responses to thermal stress are often more nuanced and species/environmentally dependent. However, symbiont ‘specialist’ corals that house *Cladocopium C15* are often more thermally resilient than others, suggesting that *C15* may also be a thermally tolerant symbiont type. What functional traits these two species carry and allow them to be more thermally tolerant than others is an active area of research both for basic and applied fields of coral conservation.

The *Symbiodiniaceae* family is genetically and phenotypically diverse, but phylogeny alone is not sufficient to explain the broad differences in ecological success that have been observed across the family (Lewis et al., 2018; Sampayo et al., 2008; Goyen et al., 2017; Suggett et al., 2017; Mansour et al., 2018). Relatively recent adaptive radiations among certain genera (Thornhill et al., 2014) driven by variable nutrient and light environments, coral skeletal architecture, and tissue pigments have presumably resulted in diverse functional traits across species, including variability in response to environmental perturbations such as thermal stress (D’Angelo & Wiedenmann, 2014; Sully & van Woesik, 2020; Scheufen et al., 2017; Suggett et al., 2015; Suggett et al., 2017). However, as there still exists a gap in knowledge regarding how genomic diversity within this group translates into functional trait differences and phenotypes across species and environments, a better understanding of algal trait variability across the *Symbiodinaceae* family is needed.

Chlorophyll-*a* fluorescence techniques are commonly employed within coral research as a tool for assessing photosynthetic health under high temperature stress (Warner et al., 1999, Rodríguez-Román et al., 2006, Cunning et al., 2021). Indeed, these tools have been a critical component of bleaching response research and are increasingly utilized to characterize trait-based differences within and across coral species (Suggett et al., 2022, Hoadley et al., 2021). Incorporation of varying light protocols and multispectral analyses (Hoadley et al., 2023, Szabó et al., 2014) further increases the utility of these tools for assessing nuanced functional trait differences across coral species and/or environmental conditions. Such tools provide the ideal platform for phenomic studies as ‘high-content’ data sets can be easily captured from individual corals and then assessed using machine learning techniques to establish photo-physiological profiles representative of key species and/or underlying cellular traits.

First-order traits, or traits which form the basis of function, are thought to be better determinants of algal survival and success than secondary traits, which arise from the performance of primary traits (Suggett et al., 2017). First-order traits such as allometric scaling (cell size) and nutrient budgeting (atomic Carbon: Nitrogen: Phosphorus ratios) likely regulate second-order traits, such as photoprotection and light utilization strategies (Suggett et al., 2017; McIlroy et al., 2020; Hoadley et al., 2021). However, first order traits are often difficult to measure, requiring destructive and sometimes expensive analytical methodology. In contrast, second order traits, such as light utilization strategies in photosynthetic organisms, are relatively easy to characterize using rapid and non-invasive tools such as chlorophyll-*a* fluorescence (Warner et al., 1999, Suggett et al., 2015, Warner & Suggett 2016). Because light utilization strategies differ across environmental conditions and species of *Symbiodiniaceae*, characterizing relationships between such secondary and underlying primary traits could provide useful insight into what drives the observed functional trait differences derived through chlorophyll-*a* fluorescence. Establishing correlations between primary traits and light utilization characteristics, future studies may also be able to infer certain primary trait characteristics via chlorophyll-*a* fluorescence alone.

In this study, we characterized photosynthetic poise of six different *Symbiodiniaceae* species living *in-hospite* among 60 Pacific coral fragments, spanning 12 coral species (*Montipora capricornis, Acropora yongei, Montipora digitata, Turbinaria reniformis, Acropora millepora, Acropora humilis, Acropora valida, Acropora* sp.*, Pavona cactus, Psammacora contigua, Pocillopora damicornis, and Cyphastrea chalcidicum*), sourced from the Reef Systems Coral Farm, Inc, in New Albany, Ohio. Prior to isolation of the algal symbionts for characterization of primary traits, we first assessed active chlorophyll-*a* fluorescence metrics using 5 excitation wavelengths during a time-resolved actinic light protocol designed to test acclimation, relaxation, and light utilization at high and low light intensities (Hoadley et al., 2023). This yielded a total of 1360 individual metrics which were then used to cluster coral colonies based on similarity of their photophysiological phenotype. Phenotypes were then compared to identified symbiont species and linked to underlying differences in primary traits across phenotypes. Next, a network analysis was utilized to elucidate specific correlations between our primary and secondary traits. Convergence between algal species and phenotypes provide a useful means for understanding trait-based differences across genetic lineages, along with the underlying cellular traits that regulate them.

## Materials and Methods

### Coral husbandry and environmental conditions

All corals were housed at Reef Systems Coral Farm (New Albany, Ohio). This facility consists of large ∼4000 L raceways (8’ x 4’ x 4’) housed within a greenhouse facility (optically clear plastic roofing with no shade cloth). Smaller indoor tanks (∼1100 L) contain additional corals under LED illumination. All tanks utilize the same artificial seawater (Reef Crystals in RO/DI filtered water) and receive frequent (10-20%, 1-2 week^-1^) water changes. All tanks contain submersible power heads (Tunze) which circulate water within each system between 50-100 times per hour. Additional overflow pumps exchange water (1-2 times volume of tank hr^-1^) with external sumps equipped with mechanical (25-micron sieve) and foam fractionation filtration. All coral fragments are mounted on ceramic disks (3cm) and attached using cyanoacrylate gel (coral glue). Peak irradiance (measured at the same tank depth for each tank) within the outdoor (greenhouse) tanks was 1050 µmol m^-2^ sec^-1^ (walz, 4pi sensor). Indoor corals were illuminated using LED lighting (Radion Xr30 Pro) on a 10:14 hr dark: light cycle with peak irradiance measured at 300 µmol m^-2^ sec^-1^. Indoor and outdoor corals thus differ both in max irradiance but also in light spectra (natural lighting vs. LED output).

Reef Systems Coral Farm (New Albany, Ohio) houses over 30 different species of coral, with thousands of individual fragments originating from coral colonies that were harvested in Fiji but have been captively grown at the facility for over ten years. All individual fragments utilized in the study had been fragmented and mounted for at least one month prior to use. In order to maximize diversity, we collected three replicate fragments from twenty different coral colonies chosen to include a wide range of life strategies and histories, including both mound and branching species acclimated to greenhouse conditions; *Psammacora contigua* (1 genotype), *Acropora yongei* (1 genotype)*, Acropora millepora* (1 genotype)*, Acropora humilis* (1 genotype)*, Acropora valida* (1 genotype)*, Acropora* species (1 genotype), *Turbinaria reniformis* (2 genotypes)*, Pavona cactus* (1 genotype)*, Montipora digitata* (1 genotype)*, Montipora capricornis* (3 genotypes)*, Cyphastrea chalcidicum* (1 genotype)*, Pocillopora damicornis* (2 genotypes). In addition, indoor acclimated fragments representing species *Acropora humilis* (1 genotype), *Turbinaria reniformis* (1 genotype), and *Montipora capricornis* (2 genotypes) were also included in our analyses.

### Chlorophyll fluorescence-based phenotyping protocol

Prior to measurements, each individual coral fragment was dark acclimated for 20-25 minutes (Suggett et al., 2015). Photophysiological responses to changing light conditions were then measured using a prototype Chlorophyll-*a* fluorometer previously described in Hoadley et al., 2023. Briefly, fluorescence induction curves were produced through excitation with 1.3-*μ*s single turnover flashlets spaced apart by 3.4 *μ*s dark intervals (32 flashlets were utilized under 420, 442, and 458-nm excitation while 40 flashlets were utilized during 505 and 520-nm excitation). Each fluorescence induction curve was followed by a 300 ms relaxation phase consisting of 1.3-*μs* light flashes spaced apart with exponentially increasing dark periods (starting with 59-*μs*). Fluorescence induction and relaxation curves measured using each excitation wavelength were sequentially repeated 5 times per measurement timepoint. These photophysiological measurements were repeated 34 times during an 11-minute variable actinic light protocol in which corals were exposed to an initial dark period (30s) followed by 3 different light intensities (300 µmol m^-2^ sec^-1^ for 3.5 minutes, then 50 µmol m^-2^ sec^-1^ for 1.5 minutes followed by 600 µmol m^-2^ sec^-1^ for 3.5 minutes), and a final dark recovery period (∼ 2 minutes) (Hoadley et al., 2023). Resulting fluorescence data was analyzed using custom R scripts (Hoadley et al., 2023) and according to equations set forth in Kolber et al., 1998 in order to calculate quantum yield of photosystem II (Φ_PSII_), PSII functional absorption cross-section (σ_PSII_), reaction center connectivity (ρ), non-photochemical quenching (NPQ), antenna bed quenching (ABQ), photochemical quenching (qP), and two time constants for electron transport (τ_1_ and τ_2_) where τ_1_ reflects acceptor-side changes of PSII and τ_2_ reflects changes in plastoquinone pool reoxidation (Suggett et al., 2022; Hoadley et al., 2023). However, downstream PQ reoxidation kinetics (τ_2_) are typically derived from a multiturnover induction and relaxation flash sequence (Suggett et al., 2022) whereas ours are derived from the second exponential component of a single turnover induction and relaxation flash sequence (τ ^ST^).

### Endosymbiont Isolation

Once chlorophyll-*a* fluorescence-based measurements were complete, a portion of coral tissue was removed using the water-pick method (Johannes & Wiebe, 1970) and filtered seawater (artificial seawater vacuum-filtered, 0.2-micron filter). The resulting tissue-water slurry was homogenized using a hand-held tissue homogenizer (tissue tearer) and then measured using a graduated cylinder. Samples were then centrifuged (8,000 X g, 10 minutes, room temperature). After centrifugation, the supernatant was removed, and the pellets were resuspended in sterile seawater, vortexed for 30 seconds and then centrifuged once more to wash the endosymbionts free of the host tissue. Resulting algal pellets were resuspended in 10ml of sterile seawater.

### Flow Cytometry (cell size, chlorophyll-*a*, neutral lipid content, and granularity)

One ml aliquots of resuspended algal samples were preserved with glutaraldehyde (0.1% final concentration), incubated in the dark for 20 minutes, flash frozen in LN_2_, and then stored at –80 C for downstream flow cytometry. All samples were analyzed using a Attune NxT (Invitrogen, USA) equipped with a 488 nm (200 mW) laser and a 200 µm nozzle. Glutaraldehyde-fixed samples were re-pelleted using a centrifuge (12,000xg, 5 minutes), resuspended in 1x filtered PBS, and then filtered through a 50-micron mesh filter to remove residual coral tissue or large cell aggregates. Samples were then diluted 4x with 1x filtered PBS and spiked with 10 µl of 1:1000 diluted fluorescent bead stock (Life Technologies 4.0 µm yellow-green 505/515 Fluospheres sulfate). 100-µl per culture was analyzed at a flow rate of 200-µl min^-1^. For characterization of *Symbiodiniaceae* cells, data collection was triggered on forward light scatter (FSC), while red (695/40 nm bandpass filter) autofluorescence detected chlorophyll-*a*, and both were utilized to gate the cell and bead populations for bead-normalized FSC, side scatter (SSC) and chlorophyll fluorescence measurements.

For quantification of neutral lipid content, 500 µl of the diluted sample was first stained with 2-µl 5mM Bodipy 493/503 (Invitrogen, 4,4-Difluoro-1,3,5,7,8-Pentamethyl-4-Bora-3a,4a-Diaza-s-Indacene) and then incubated in the dark at 37LJC for 15 minutes. Stained samples were then run according to the same conditions described above. The *Symbiodiniaceae* cell population was identified using FSC-height and autofluorescence, and the gated population’s green fluorescence was quantified (bandpass filter 530/30).

### Total C:N:P Content and Nutrient Analysis

For Carbon and Nitrogen analysis, 2-ml of each resuspended algal sample was filtered through a 13mm ashed GF/F filter and dried in a 95LJC oven for 24 hours. Filters were packed into tin capsules, combusted (Costech Instruments 4010 Elemental Combustion System) and analyzed via Elemental Analyzer. Total carbon and nitrogen values were compared to an atropine standard. For particulate organic phosphorus (POP) analysis, 2-ml of each isolate was filtered through a 13mm ashed GF/F filter and stabilized with a 0. 17M N_a_2SO_4_ rinse. Filters were placed in muffled scintillation vials with 2-ml aliquots of 0.017M N_a_2SO_4_ and evaporated to dryness in a 95□C muffle oven for 24 hours. Vials were covered in aluminum foil and baked at 450□C for 2 hours, and then were baked with 5-ml 0.2 M HCl at 90□C for 30 minutes. Samples were diluted with 5-ml ultra-pure water and analyzed using the Skalar SAN+ system and compared to an Adenosine Triphosphate standard.

To confirm that corals across different tanks were grown under similar nutrient conditions, 20-ml samples from each husbandry tank were collected and stored at –20□C until analysis, when they were thawed and loaded into the Skalar SAN+ system’s autosampler. The samples were analyzed for μM nitrate, nitrite, ammonia, and phosphate via continuous flow analysis and according to EPA standard techniques (EPA 1993a, EPA 1993b).

### DNA Sequencing

A 2-ml aliquot of isolated symbiont cells were first pelleted (centrifugation at 10000 rcf for 5 minutes) and then stored in 1-ml of DMSO buffer solution (Pettay et al., 2015) at 4□C. DNA was extracted from each sample using the Wizard Genomic DNA Purification Kit (Promega). Quality of DNA samples were assessed on a 1.0 Nanodrop (Thermo Scientific) and all samples had 260/280 and 260/230 values above 1.5. For each sample, PCR was first performed targeting the internal transcribed spacer 2 (ITS2) region using the previously established primer pairs (ITSintfor2: 5’GAATTGCAGA ACTCCGTG-3’, ITS2-reverse: 5’GGGATCCATA TGCTTAAGTT CAGCGGGT-3’) (LaJeunesse et al., 2002; Sheffield et al., 1989). Resulting amplicons were subjected to a second round of PCR (only 8 cycles) using the same primer pairs with adapter sequence (Forward-<u>TCGTCGGCAGCGTCAGATGTGTATA</u> <u>AGAGACAG</u>GAATTGCAGAACTCCGTG; Reverse-<u>GTCTCGTGGGCTCGGAGATGTGT ATAAGAGACAG</u>GGGATCCATATGCTTAAGTTCAGCGGGT). Adapter sequences are underlined in the above primer sets. All PCR was achieved using Hot Start Taq DNA Polymerase (New England BioLabs, Inc) under the following settings: denaturation 94.0LJC for 30s, annealing 54.0LJC for 35s, extension 68.0LJC for 3min x 35 cycles (8 cycles in round two), final extension 68.0LJC for 5 minutes. Samples were then purified using the GeneJET PCR Purification Kit (Thermo Scientific) and visualized on agar gels to confirm results. Amplicons were submitted to the University of Delaware Sequencing and Genotyping Center for library preparation and metagenomic sequencing. Amplicons were dual indexed using the Nextera system and were run as a single library using a paired-end 300 base pair x 2 strategy on an Illumina MiSeq system. Demultiplexed FASTQ sequences were then uploaded to SymPortal for profiling (Hume et al., 2019). Given known high variability in rDNA copy numbers across species and genera (LaJeunesse & Thornhill, 2011) which may hinder translation of data into accurate relative abundances (Davies et al., 2022), absolute abundance ITS2-type profiles were then normalized to rDNA copy numbers according to Saad et al., 2020.

### Statistical Analyses

All statistical analyses were conducted in R (version 4.1.3). Algal genotypes for each coral sample were derived from SymPortal-generated ITS2 profiles normalized to reflect relative abundance of each symbiont genera. Importantly, known differences in *Cladocopium* and *Durusdinium* ITS2 copy numbers was addressed by normalizing to previously derived rDNA copy ratios of 2119:362 (Saad et al., 2020). The dominant (> 70% of ITS2 sequences) symbiont type in each coral fragment (Fig. 2b) was then utilized to screen for photophysiological metrics that were significantly different across algal species using either a one-way ANOVA or Kruskal-Wallis test if data did not meet the assumptions of normality. The resulting dataset was then transformed using Z-scores and then plotted as a heatmap (Fig. 2a) with individual coral colonies represented by each column, and photophysiological metrics across rows. A clustering analysis (1000 bootstrap iterations) was performed using the R packages pvclust (Suzuki & Shimodaira, 2006) and dendextend (Galili, 2015) to cluster individual coral samples by algal phenotype. Clustering analyses were also carried out on heatmap rows and resulting clusters were further analyzed using custom scripts to identify which photophysiological metrics were most important for separating our coral fragments into separate phenotypes.

**Figure 1.**
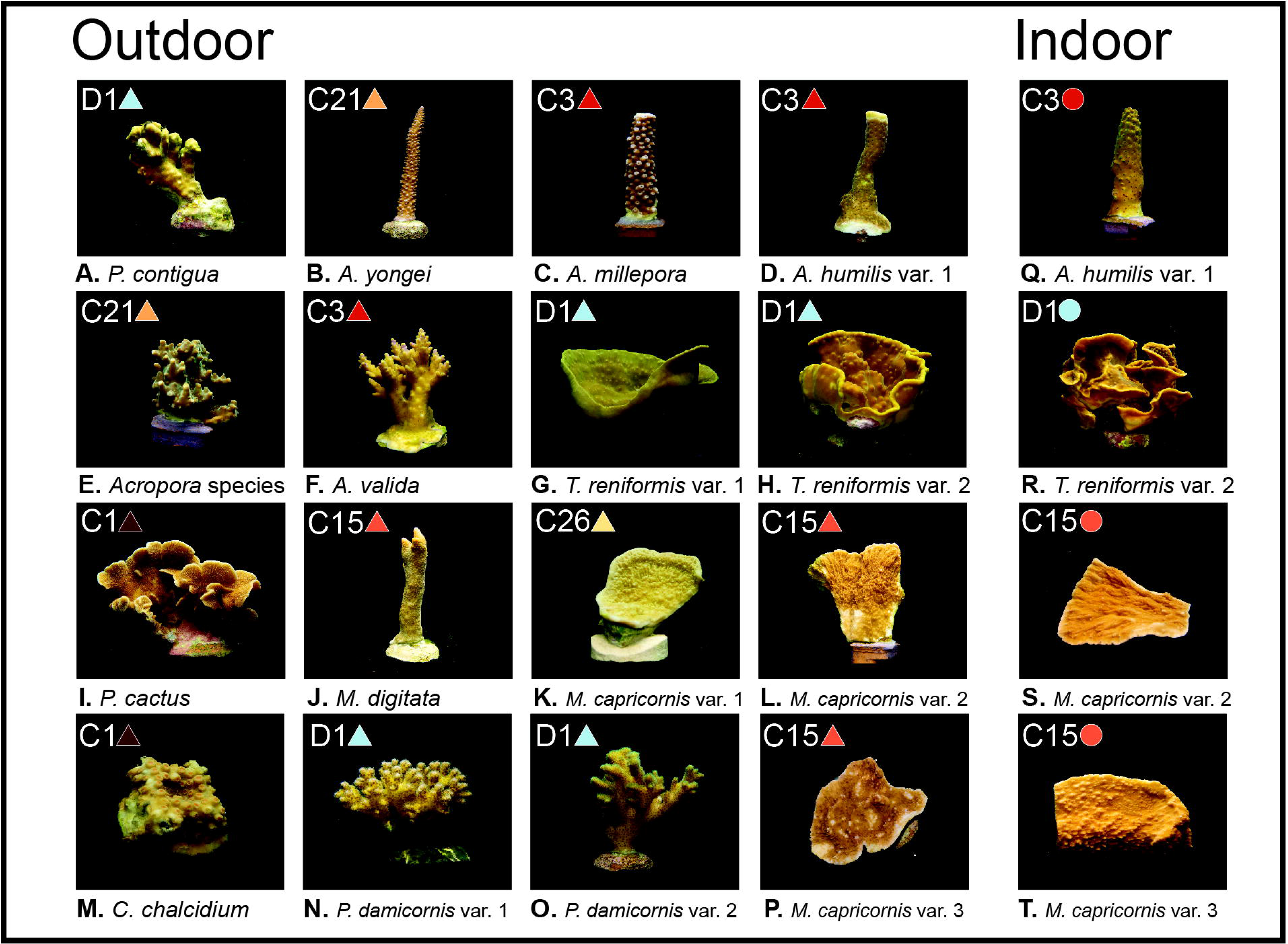
– Coral images, light environments, and dominant symbiont types. Panels a-t show the 12 coral species with 1-3 variants per species, separated by growth environment (Outdoor grown corals on the left and indoor grown corals on the right). Dominant symbiont type found for each coral is included in the top left of each panel, along with corresponding colored symbols (circles and triangles) which are utilized throughout the remaining figures to identify symbiont type. All photos taken by Audrey McQuagge.

**Figure 2.**
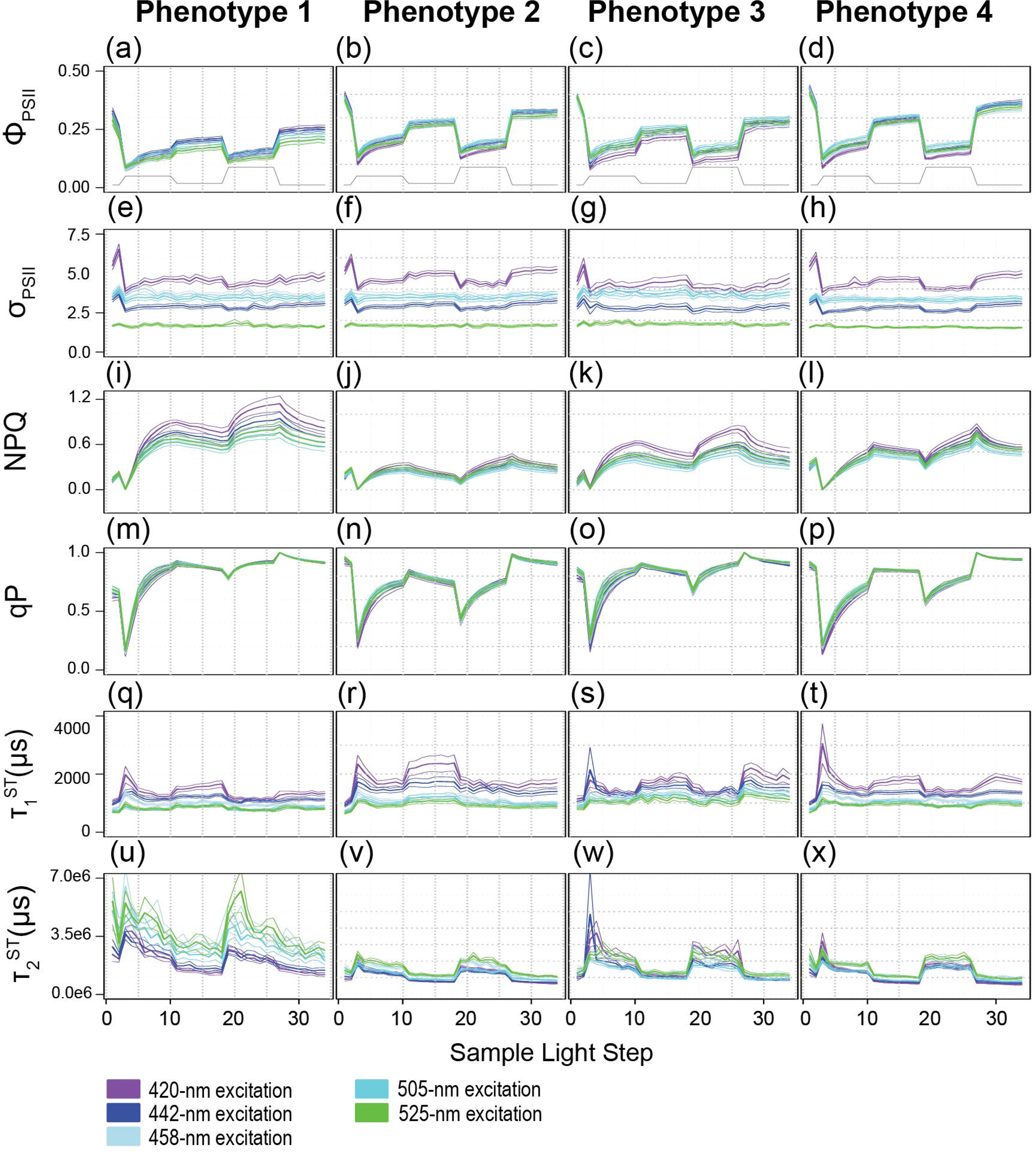
– Heat map with dendrogram and relative abundance bar graphs. The heatmap (a) analysis reflects a total of 987 photophysiological biometrics which were found to differ significantly across symbiont types. Dominant photo-physiological metrics within each of the four identified row clusters are displayed on the right of the heatmap. The colored dendrogram above the heat reflects 4 distinct phenotypes with resulting bootstrap support indicated at each major branch. The larger bar graph directly below the heat map (b) represents the relative abundance of symbionts within each coral sample whereas the second (c) and third (d) bar graphs represent host coral genera and coral growth environment (indoors or outdoors) respectively. Capital letters underneath the bar graphs represent the coral species listed under the various panels in Fig. 1. Letters that are in bold indicate that all three fragments for that coral colony are found in the same phenotype.

Full actinic light profiles for identified photophysiological metrics (Fig. 2a) were plotted in Fig. 3 and a repeated measures linear mixed model with a tukey posthoc (with Bonferroni correction) identified significant differences across algal phenotypes (Supplemental Table 4) using the lmerTest (Kuznetsova et al., 2017) and multicomp (Hothorn et al., 2008) R packages. Spectrally-dependent differences within each photophysiological metric and algal phenotype were similarly assessed (Supplemental Table 3). Significant differences in cellular physiology (cell size, Granularity, Chl *a*, N:P, C:P, and C:N ratios) across phenotypes was also assessed. For each metric, normality was first determined (Shapiro-Wilks). If data were determined to be normal, a One Way ANOVA followed by a Tukey posthoc was performed. For data that did not meet the assumptions of normality, a Kruskal Wallace with Bonferroni correction was performed (Fig. 4).

**Figure 3.**
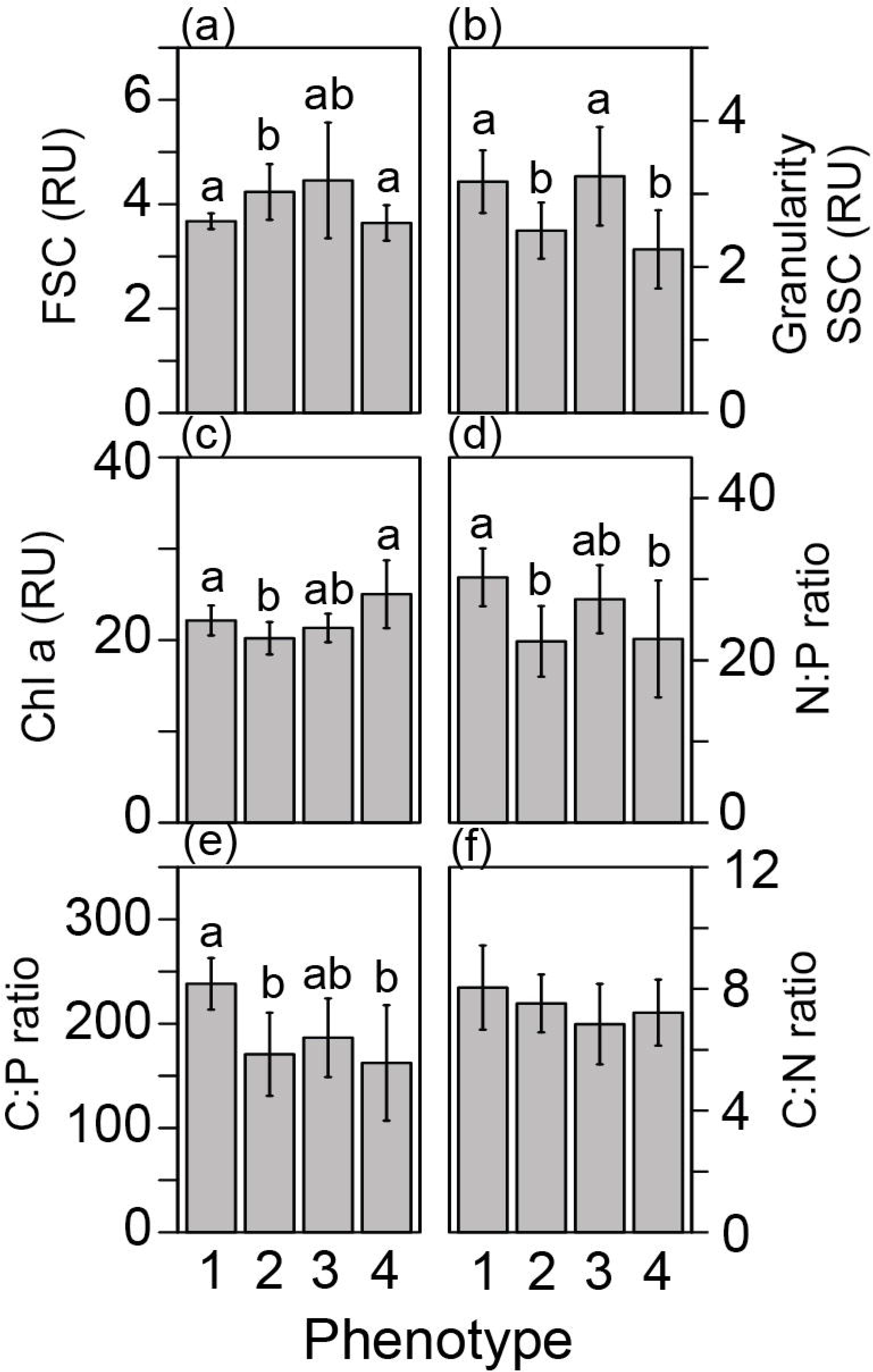
– Profiles for photophysiological biometrics driving variability across phenotypes. Average (± standard error) traces for photo-physiological metrics identified in Fig. 2 as contributing significantly towards establishing the four phenotypes across our coral colonies. Phenotypes 1-4 are displayed from left to right. Panels **a-d** reflect the Quantum Yield of PSII (Φ_PSII_), **e-h** reflect the absorption cross-section of PSII (σ_PSII_), **i-l** reflect non-photochemical quenching (NPQ), **m-p** reflect photochemical quenching (qP), **q-t** and **u-x** reflect the reoxidation constants τ_1_ ^ST^ and τ ^ST^ respectively. Line color indicates excitation wavelength with purple representing 420-nm; dark blue, 442-nm; light blue, 458-nm; teal blue, 505-nm, and green, 525-nm. The grey line on panels a-d displays the variable light protocol. Bonferroni-adjusted p-values for comparisons across excitation wavelength and phenotype can be found in Supplemental Tables 3 and 4.

**Figure 4.**
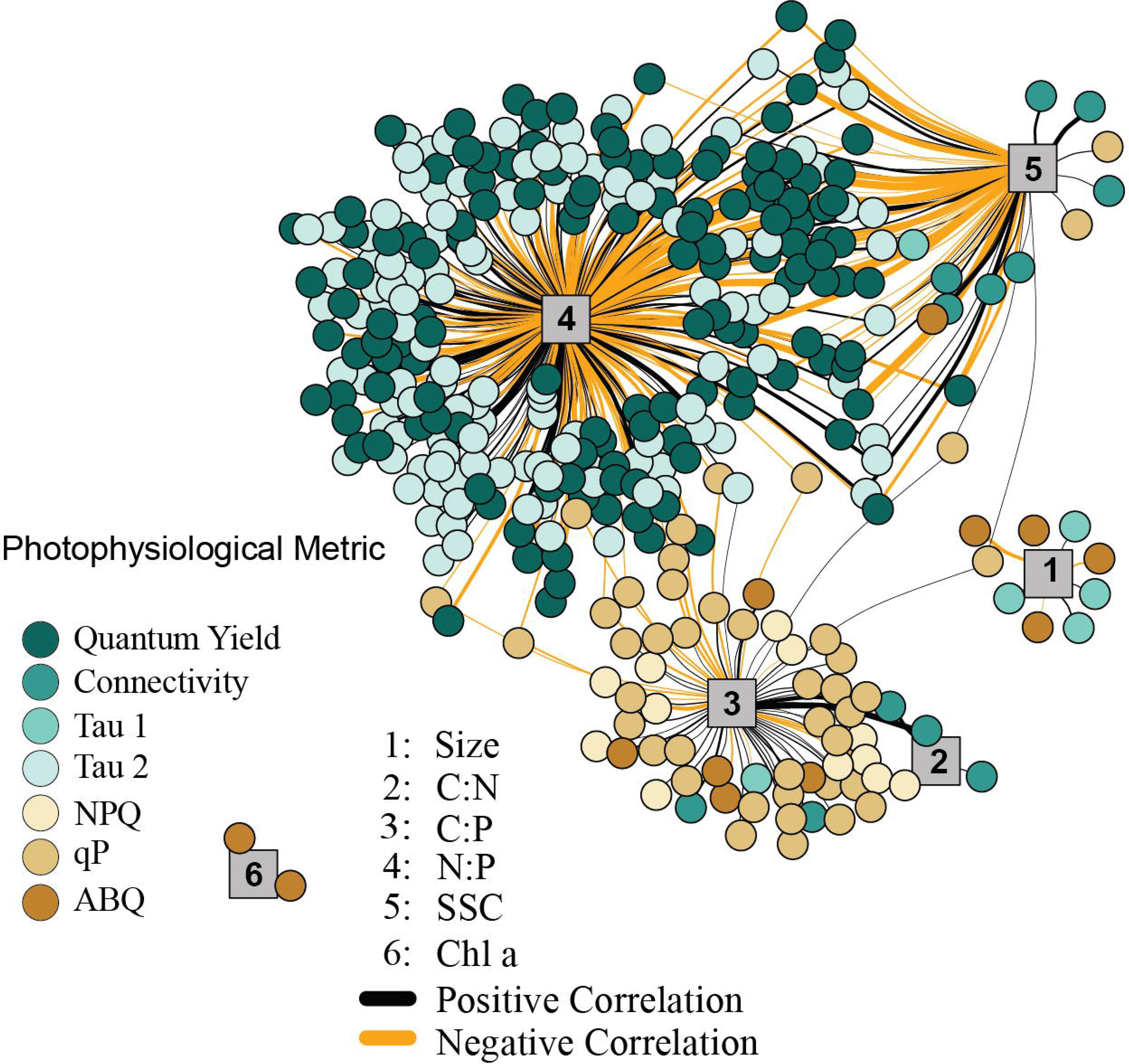
– Differences in cellular physiology across phenotypes. Differences in the underlying symbiont cellular physiology was compared across our four photo-physiologically derived phenotypes. Flow cytometrically derived cell size (**a)** granularity (**b**), chlorophyll-*a* fluorescence (**c**), Nitrogen to phosphorous (**d**), Carbon to phosphorous (**e**), and Carbon to nitrogen (**f**) ratios are represented as the mean (± standard error) for each phenotype. Different letters above the bars in each panel reflect significant differences (Tukey posthoc) across phenotypes.

A network analysis was employed to look for significant correlation between photophysiological metrics and primary cellular traits using averaged values across coral species replicates (3 species^-1^). Only correlations with a Pearson value above 0.6 were utilized. Using the igraph package (Csardi & Nepusz, 2006), specific correlations (Pearson’s value of 0.55 or above) between symbiont cellular metrics and fluorescence-based phenotyping data were identified and are displayed in Fig. 5.

**Figure 5.**
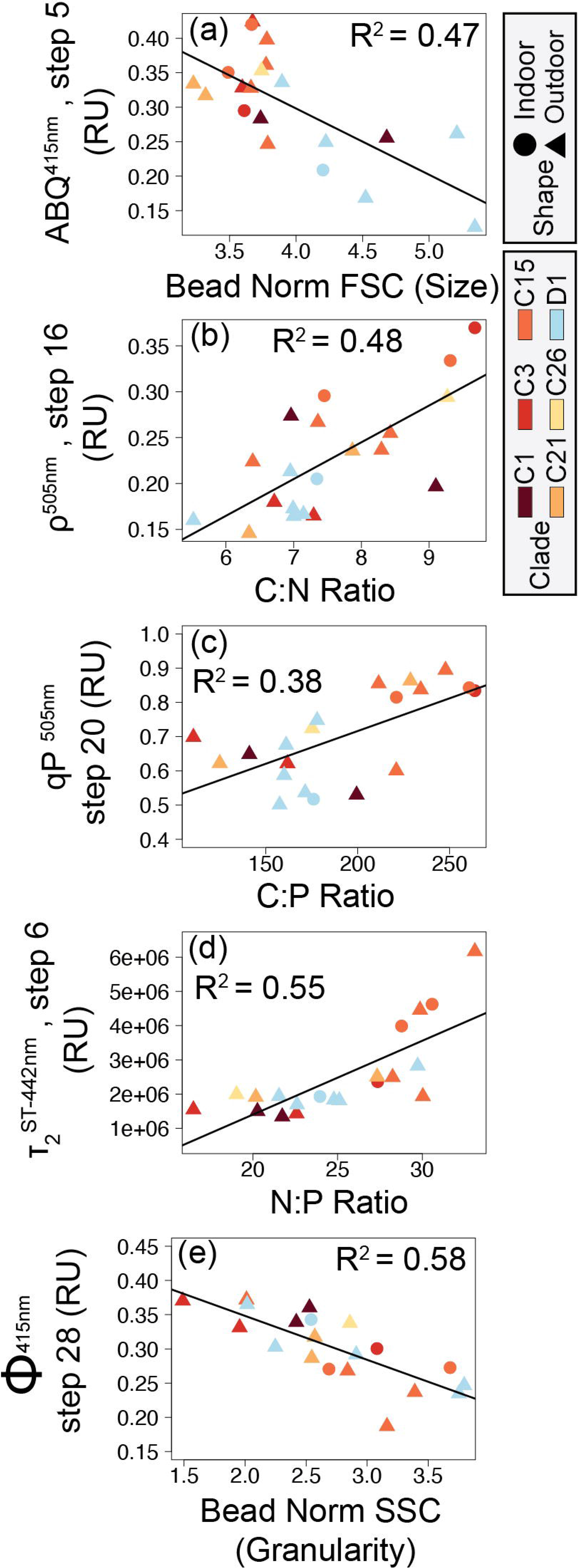
– Network analysis between cellular and photophysiological traits. The network analysis only reflects significant correlations between first (cellular) and second (photo–physiological) order traits. Cellular traits are indicated by numbered grey vertices, **1** = cell size, **2** = C:N, **3** = C:P, **4 =** N:P, **5 =** Granularity, **6 =** Chl *a*. Line thickness corresponds to the strength of the correlation (between 0.6 and 0.9 Pearson R values), with thicker lines representing traits that are more highly correlated. Positive correlations are indicated by black lines, while negative correlations are indicated by orange lines. Colored circles reflect the various photophysiological variables with strong (> 0.6 Pearson R) correlation to underlying cellular metrics.

## Results

### Symbiont types

Symportal analysis revealed that most corals were dominated by a single genotype (>70% relative abundance) of *Symbiodiniaceae*. *Acropora* (including *A. yongei*, *A. millepora*, *A. humilis*, *A. valida*, and unknown species) predominantly hosted either *C3* or *C21* symbiont types, while *Montipora* (including *M. capricornis* and *M. digitata*) hosted primarily *C15*, but sometimes *C26*, variants (Fig. 2b). The symbiont type *D1* was primarily observed in *Turbinaria*, *Psammacora*, and *Pocillopora*.

### Phenotype to genotype clustering and profiles

Of the 1360 algal biometrics derived from the fluorescence-based excitation profile, 987 were found to be significantly (*p<*0.05) different across the dominant symbiont species (*C1*, *C3*, *C15*, *C21*, *C26*, and *D1*, Table 1). These identified algal biometrics were then utilized to organize samples according to trait-based phenotypes (Fig. 2a), with the resulting dendrogram organized into four distinct phenotypes according to the largest clustering groups (Fig. 2a). Phenotype 1 contains 14 of the 15 *C15-*dominated coral colonies (*Montipora digitata* and *Montipora capricornis*), with both high and low-light acclimated fragments clustering together (Fig. 2). Two additional fragments of *Acropora sp.* (*C3*) were also found in phenotype 1. Phenotype 2 was predominantly (12 of 16 fragments) comprised of *Durusdinium trenchii* (*D1)-*dominated corals (low-light *Turbinaria reniformis, Pocillopora damicormis,* and *Psammacora contigua*). The remaining four coral fragments in phenotype 2, were dominated by *Cladocopium C1* (in *Cyphastrea chalcidicum* or *Pavona cactus*). Coral fragments belonging to phenotype 3 were comprised of 3 *A. millepora* fragments (*C3*) and 5 fragments of high-light acclimated *Durusdinium D1-*dominated *Turbinaria renifomis*. Lastly, phenotype 4 was comprised of both high and low-light acclimated *A. humilis* (*C3*) fragments, along with all *C21* dominated corals (*A. yongei* and *Acropora sp.*), all three fragments of *M. digitata* dominated by *C26*, two fragments of *P. cactus* (*C1*) and single *C15-*dominated (*M. digitata*) and *D1*-dominated (*T. reniformis* – high light acclimated) fragments. Of the 20 coral colonies represented in this study, only four contained fragments which did not all cluster within the same phenotype. Based on row clustering and custom scripts, quantum yield of PSII (Φ_PSII_), functional absorption cross section of PSII (σ_PSII_), non-photochemical quenching (NPQ), photochemical quenching (qP), along with the reoxidation kinetics (τ_1_ ^ST^ and τ ^ST^) were determined to be the primary drivers for the observed phenotypic structure and were thus plotted in full detail (Fig. 3) for each of the four phenotypes described above.

**Table 1:** Coral hosts, growth environments, and relative abundances of ITS2 symbiont types determined through SymPortal.

| Coral Host | Growth Environment | Dominant Symbiont Type (full ITS2 Name) | Rel. Abundance | Secondary Symbiont Type (Full ITS2 Name) | Rel. Abundance |
| --- | --- | --- | --- | --- | --- |
| *A. humilis* var 1 | Outdoor | C3k/C3-C50a-C3dq-C50f-C3ba-C3a | 1.00 |  |  |
| *A. millepora* | Outdoor | C3/C3u-C115-C21ab-C3ge | 1.00 |  |  |
| *Acropora* sp. | Outdoor | C21-C3-C21ag-C21af | 1.00 |  |  |
| *A. valida* | Outdoor | C3z-C3-C3.10-C115-C3an | 0.98 | C21 | 0.02 |
| *A. yongei* | Outdoor | C21-C21ag-C3-C21as | 1.00 |  |  |
| *C. chalcidium* | Outdoor | C1/C3-C1c-C1b-C42.2-C1bh-C1br | 1.00 |  |  |
| *M. capricornis* var 1 | Outdoor | C26A-C21 | 0.98 | C1 | 0.02 |
| *M. capricornis* var 2 | Outdoor | C15-C15he-C15ed-C15cq-C15vl | 1.00 |  |  |
| *M. capricornis* var 3 | Outdoor | C15-C15he-C15ed-C15cq-C15vl | 1.00 |  |  |
| *M. digitata* | Outdoor | C15/C15gb-C15vk | 1.00 |  |  |
| *P. cactus* | Outdoor | C1b/C1/C1mm-C3-C1u-C1dg | 1.00 |  |  |
| *P. contigua* | Outdoor | D1-D4-D4c-D1h | 0.98 | C1ec/C1-C1b-C3 | 0.02 |
| *P. damicornis* var 1 | Outdoor | D1/D6-D1h-D1kg-D1ke-D1kf-D1kh | 0.98 | C1d/C1/C42.2/C3-C1b-C3cg-C45c-C115k-C1au-C41p | 0.02 |
| *P. damicornis* var 2 | Outdoor | D1/D6-D1h-D1kg-D1ke-D1kf-D1kh | 0.84 | C1d/C1/C42.2/C3-C1b-C3cg-C45c-C115k-C1au-C41p | 0.16 |
| *T. reniformis* var 1 | Outdoor | D1-D4-D1ki-D1c | 1.00 |  |  |
| *T. reniformis* var 2 | Outdoor | D1-D4-D4c-D1c-D2-D1k | 0.99 | C3z-C3-C3.10-C115-C3an | 0.01 |
| *A. humilis* var 2 | Indoor | C3k/C3-C50a-C21ab-C50f-C3ba-C3dq | 0.97 | C1/C1c | 0.03 |
| *M. capricornis* var 2 | Indoor | C15-C15he-C15ed-C15cq-C15vl | 0.95 | D1/D6-D1h-D1kg-D1ke-D1kf-D1kh | 0.05 |
| *M. capricornis* var 3 | Indoor | C15-C15he-C15ed-C15cq-C15vl | 0.98 | D1/D6-D1h-D1kg-D1ke-D1kf-D1kh | 0.02 |
| *T. reniformis* var 2 | Indoor | D1-D4-D1ki-D1c | 0.97 | C1/C21/C3-C1b-C1c-C42.2-C1bh | 0.03 |

A mixed linear model was utilized to identify spectrally dependent differences across phenotypes for the photo physiological metrics, Φ_PSII_, σ_PSII_, NPQ, qP, τ_1_ ^ST^, and τ ^ST^ reoxidation kinetics (Fig. 3 and Supplemental Table 4). For all excitation wavelengths except 420nm, phenotype 1 had significantly (*p <* 0.0159) lower Φ_PSII_ values compared to all other phenotypes. Under 420nm excitation, Φ_PSII_ profiles for phenotype 1 were significantly (*p <* 0.0001) lower than phenotypes 2 and 4, but not phenotype 3. Additionally, Φ_PSII_ profiles for phenotype 3 were also significantly (*p <* 0.001) lower than phenotype 2 but only under 420nm and 442nm excitation whereas phenotype 3 different significantly (*p <* 0.043) from phenotype 4 under all excitation wavelengths except 525nm. No differences across phenotypes were observed for σ_PSII_ under any excitation wavelength and indicate that subtle differences may only exist when comparing across specific symbiont species (not phenotypes). Under all excitation wavelengths, nonphotochemical quenching profiles reached the highest values in phenotype 1 whereas relatively small changes were observed in phenotype 2. For phenotypes 3 and 4, NPQ values did not differ significantly from one another but represent a medium level that is significantly (*p* < 0.001) different from phenotypes 1 and 2. Photochemical quenching (qP) profiles observed in phenotypes 1 and 3 differed significantly (*p* < 0.003) from those in phenotype 2 and 4 under all excitation wavelengths. Differences in τ ^ST^ were more sporadic across phenotypes as profiles under 420nm excitation differed significantly (*p* = 0.001) between phenotypes 1 and 2. Under 442nm excitation, τ ^ST^ profiles for phenotype 1 were significantly (*p <* 0.011) different from those observed for phenotype 2 and 3. For τ_1_^ST^ profiles measured under 505nm excitation, phenotype 1 was significantly (*p <* 0.021) different from all others, while phenotypes 3 and 4 were also differed (*p* < 0.027) from one another. Lastly, τ_2_^ST^ profiles for phenotype 1 had significantly (*p* < 0.002) slower (higher time constants) kinetics than those observed for all other phenotypes under all excitation wavelengths except 420nm. Under 420nm excitation, τ_2_ ^ST^ profiles for phenotypes 1 and 3 were only significantly (*p* < 0.002) elevated over those found in phenotypes 2 and 4.

A mixed linear model was also utilized to compare spectrally dependent photo-physiological profiles within each phenotype (Fig. 3 and Supplemental Table 3). For Phenotype 1, Φ ^420^ Φ ^442^ and Φ ^505^ profiles were significantly (*p <* 0.004) higher than Φ ^458^ and Φ ^525^. For phenotype 2, Φ ^420^, Φ ^458^, and Φ ^525^ profiles were lower (*p <* 0.005) than Φ ^442^ and Φ ^505^. For phenotype 3, Φ ^420^ appeared to be significantly (*p* < 0.006) lower than all other profiles whereas Φ ^505^ was significantly (*p* < 0.007) higher than the rest. For phenotype 4, Φ ^420^ and Φ ^458^ profiles were on average lower (*p* < 0.029) than Φ ^442^ and Φ ^505^ profiles. For all four phenotypes, spectrally dependent σ differed significantly (*p* < 0.021) from one another with σ ^420^ showing the highest and σ ^525^ the lowest values overall. Interestingly, and in contrast to that observed for σ_PSII_, no spectrally dependent differences in qP were observed within any phenotype. NPQ^420^ and NPQ^442^ displayed significantly (*p* < 0.0001) higher values as compared to NPQ^458^, NPQ^505^, and NPQ^525^ profiles within phenotype 1. For phenotype 2, NPQ^458^ profiles were similar to NPQ^442^ and NPQ^505^ whereas all others differed significantly (*p* < 0.001) from one another, as NPQ^420^ values tended to be slightly higher than the rest. For phenotypes 3 and 4, the NPQ^420^ profile generated significantly (*p* < 0.001) higher values whereas NPQ^505^ values were significantly (*p* < 0.034) lower than all others. All τ_1_^ST^ profiles for phenotypes 2 and 3 displayed significantly different responses from one another whereas τ ^505^ and τ ^525^ were similar to one another within phenotypes 1 and 4. Overall, τ_1_ ^420^ and τ ^442^ produces slower reoxidation kinetics as compared to τ_1_ ^458^, τ_1_ ^505^, and τ_1_ ^525^. Lastly, τ_2_ ^420^ and τ ^442^ values were significantly lower than all others in phenotypes 1 and 2. For phenotype 3, τ_2_ ^442^ and τ_2_ ^505^ produced significantly (*p <* 0.004) lower values than τ_2_ ^420^, τ_2_ ^458^, and τ_2_ ^425^. In contrast, τ_2_ ^420^, τ_2_ ^442^ and τ_2_ ^505^ profiles produced significantly (*p <* 0.001) lower values than those observed for τ ^458^, and τ ^525^.

Underlying differences in cellular physiology were also compared across the 4 fluorescence-based phenotypes (Fig. 4). Cell size (FSC) was significantly (*p <* 0.002) higher in phenotype 2 as compared with phenotypes 1 and 4 (Fig. 4a). Granularity (SSC) was significantly (*p <* 0.008) higher in phenotypes 2 and 4 as compared to phenotypes 1 and 3 and may indicate differences in light scattering abilities across groups (Fig. 4b). Fluorescence-based chlorophyll-*a* measurements were significantly (*p* < 0.025) lower in phenotype 2 as compared with phenotypes 1 and 4 (Fig. 4c). N:P and C:P ratios were significantly (*p* < 0.004) higher in phenotype 1 as compared with phenotypes 2 and 4 (Fig. 4d-e).

### Network analysis and Correlation Plots

In order to look for broad connections between primary (cellular) and secondary (photophysiological) traits, a network analysis (Fig. 5) was used to search for significant correlation between each of the 1360 fluorescence-based measurements and traditional cellular characteristics (Carbon per cell, Nitrogen per cell, Phosphate per cell, C:N ratio, N:P ratio, Cell Size, Chlorophyll-*a* (FSC), Granularity (SSC), and neutral lipids). Our analysis identified 415 correlations having a significant Pearson value of 0.6 or above. The cellular metrics N:P, C:P, and SSC displayed the greatest number of significant correlations (269, 63, and 70 respectively). For N:P ratios, the majority of positive correlations were with τ ^ST^ measurements, while most negative correlations were with Φ_PSII_ values. For C:P ratios, most correlations occurred with NPQ or qP values while SSC correlated more broadly with various metrics including Φ_PSII_, τ ^ST^, qP and connectivity. A select number of these cellular to photo-physiological correlations are displayed in full detail in Fig. 6.

**Figure 6.**
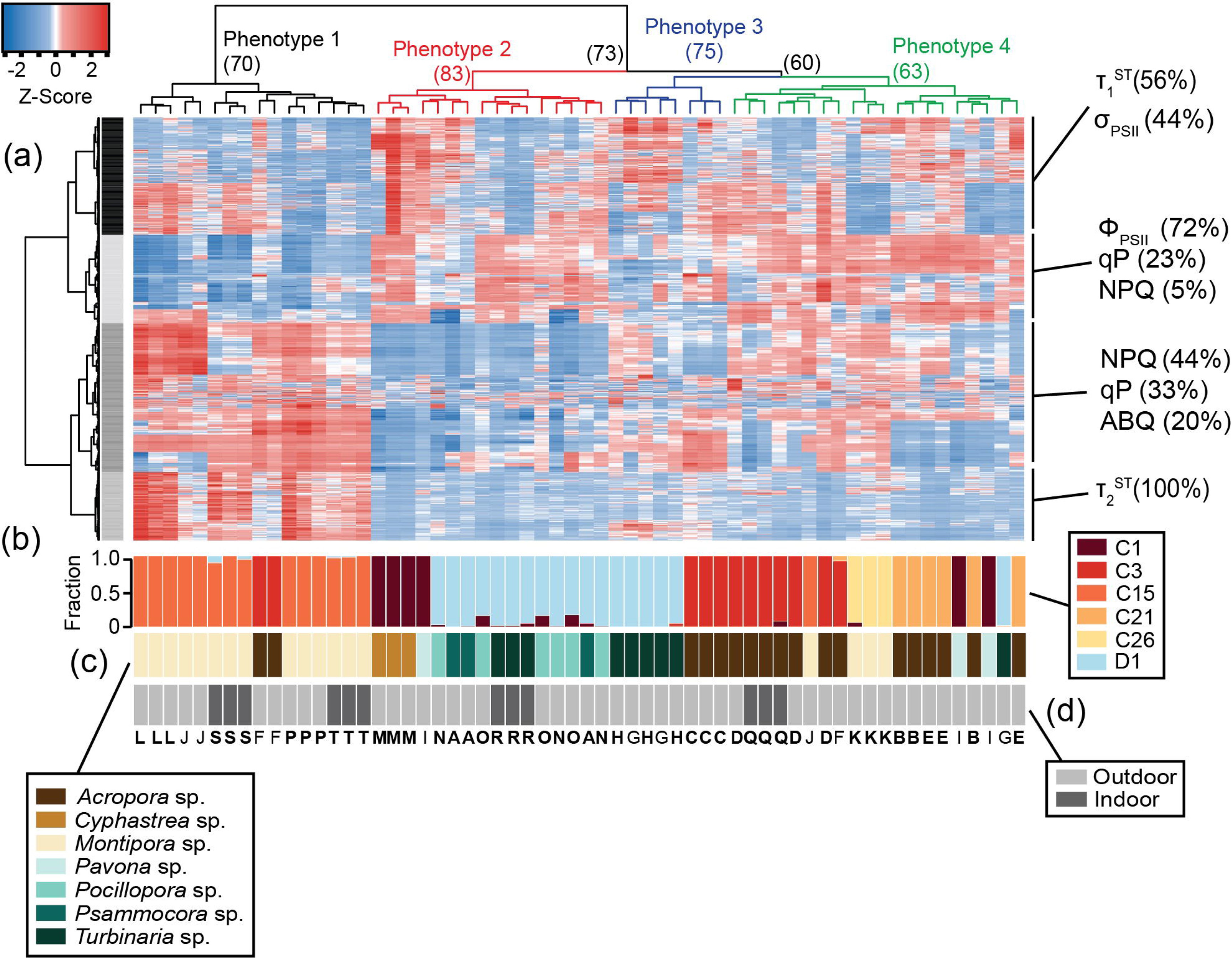
– Correlation plots between symbiont cellular and photophysiological parameters. Five correlations between cellular and photo-physiological traits with high Pearson R^2^ values are displayed in panels (**a**) ABQ vs. Cell size, (**b**) Connectivity vs. C:N ratio, (**c**) qP vs. C:P ratio, (**d**) τ ^ST^ vs. N:P ratio, and (**e**) Φ_PSII_ vs. Granularity. Point shapes indicate coral growth environment (triangles: outdoor, circles: indoor) while point color indicates ITS2 symbiont type.

## Discussion

Molecular and physiological techniques are commonly utilized by the coral research community to better understand what underpins genetic diversity and the broad range of environmental tolerances observed within the *Symbiodiniaceae* family. While the two *Symbiodiniaceae* genera *Cladocopium* and *Durusdinium* are separated by over 100 million years of evolutionary history (LaJeunesse et al., 2018), fluorescence-based phenotypes from our analysis did not entirely converge across these broad genetic designations spread across our 12 coral colonies reared under high and low light conditions (Fig. 1). For example, phenotypes 2 and 3 are comprised of corals with both symbiont genera, indicating high functional trait similarity despite large genetic differences (Fig. 2a). In contrast, greater phenotypic disparity is noted across some of the five *Cladocopium* species in this study and may reflect the relatively high genetic diversity observed in this genera as compared to others (Thornhill et al., 2014; LaJeunesse et al., 2018). Nevertheless, the degree of phenotype to genotype convergence observed within our heatmap analysis is notable, especially within the context of potentially contributing sources of functional trait disparity such as host species and light environment. High content chlorophyll-*a* flourescence-based phenotyping is already proving useful for understanding functional trait differences and their application to ecosystem services (Suggett et al., 2022, Hoadley et al., 2023), and this study further showcases the technique’s utility even across environmental light gradients while also providing direct links with underlying cellular physiology which likely regulate the observed photo physiological traits.

### ‘High content’ chlorophyl-a based phenotypes

Light acclimation state can mask species-specific differences in certain physiological metrics, as higher irradiances often lead to upregulation of stress-mitigating pathways (Ragni et al., 2010) and a reduction in photopigment production (Hoadley & Warner, 2017). This has traditionally made it difficult to capture species-specific trait-based differences without first accounting for light acclimation state. However, differences in the degree of impact that light acclimation state has on photophysiology may be largely species dependent, as high– and low-light acclimated fragments of *Cladocopium C15 and C1* (in *A. humilis*) all clustered within genotypes 1 and 4 respectively, indicating minimal impact of light acclimation state on the overall photophysiological phenotype derived through our ‘high content’ chlorophyll-*a* fluorescence protocol (Fig. 2). In contrast, high– and low-light acclimated *T. reniformis* coral fragments containing *Durusdinium D1* symbionts clustered into separate phenotypes, as did two of the low-light acclimated *C3*-dominated *Acropora humilis* coral fragments (Fig. 2). Suggett et al., 2022 also noted that the variance in light acclimation state (light niche plasticity) differed across three different species of coral found along the same reef system and at a similar depth. Understanding how various environmental factors constrain individual coral and symbiont species combinations and their underlying phenotypes will become increasingly important, especially as trait-based approaches are further applied toward coral restoration and conservation practices (Voolstra et al., 2021a).

While the degree of thermal tolerance can vary across specific host/symbiont combinations (Suggett et al., 2017), comparatively high bleaching resistance in *Durusdinium D1* and *Cladocopium C15* has been a focal point of coral research (Voolstra et al., 2021b). Whether bleaching resistance is derived through similar functional traits or if mechanisms of thermal tolerance differ across the two species is currently unknown. Coral endosymbiont thermal tolerance is often linked to photochemistry (Fitt et al., 2001; Wang et al., 2012; Warner et al., 1999), yet phenotypes differed across *Cladocopium C15* and *Durusdinium D1* dominated coral fragments in this study (Fig.s 2-3), suggesting differences in their photosynthetic poise. For example, higher reliance on non-photochemical quenching (NPQ) in response to rapid changes in light is noted for the *Cladocopium C15-*dominated phenotype (phenotype 1 – Fig. 3i) whereas *Durusdinium D1* symbionts from phenotype 2 and 3 relied more heavily on photochemical quenching to mitigate excess excitation energy as a larger proportion of PSII reaction centers remain closed throughout the actinic light protocol (Fig. 3n-o). Reoxidation kinetics between the *C15* and *D1* phenotypes also differed as phenotype 1 had lower τ_1_^ST^ values than phenotype 2 (under 420, 442, and 505nm excitation) or 3 (under 442, 458, 505, and 525nm excitation) and indicate faster rates of electron transport between the Q_a_ and Q_b_ sites within the PSII reaction centers of *Cladocopium C15* symbionts (Fig. 3q-s). Interestingly, faster τ ^ST^ kinetics for C15 were coupled with much slower τ_2_^ST^ rates as compared to other phenotypes (Fig. 3u-w). Importantly, τ ^ST^ values are derived from a 2 exponential equation fit model and thus do not necessarily reflect a specific rate constant (Hoadley et al., 2023), the higher values likely indicate slower rates of electron transport within and downstream of the plastoquinone pool. Alternative electron sinks or cyclic electron transport can play an important role in coping with excess excitation energy (Roberty et al., 2014; Vega de Luna et al., 2020) and differences in their utility across symbiont species may help drive the different reoxidation profiles observed here. Overall, stark contrasts in how each species copes with light energy during rapid changes in light are perhaps not surprising given the > 100 million years of evolutionary history that separate the two species (LaJeunesse et al., 2018). How and if these different functional traits drive unique thermal acclimation strategies will need to be the focus of a future study.

Use of multiple LED colors to excite chlorophyll*-a* fluorescence allows for potential differences in photopigment utilization to be incorporated into our phenomic analysis. For example, spectrally-dependent variance in NPQ responses may point towards differences in how photopigments are utilized to cope with excess excitation energy within phenotypes 1 and 3 (Fig. 3i, k). In contrast, phenotype 2 displayed much lower levels of NPQ in response to changes in actinic light, and little spectral variance in its profile (Fig. 3j). Non-photochemical quenching broadly encompasses various mechanisms utilized by photosynthetic organisms to dissipate harmful excess excitation energy absorbed by light harvesting antennae (Lacour et al., 2020). For many eukaryotic photoautotrophic taxa, the xanthophyll cycle (XC) is a major energy dissipation mechanism regulating observed changes in NPQ. While XC is not always involved in NPQ regulation, and its role in the family *Symbiodiniaceae* is not fully resolved, excess light energy is dissipated as heat through the inter-conversion of the photopigments (zeaxanthin to violaxanthin) or (diadinoxanthin to diatoxanthin). The sum of these various photopigments are collectively known as the xanthophyll pool, and higher concentrations are often associated with acclimation to high light (Lacour et al., 2020; Schuback et al., 2021). Importantly, these different photopigments have unique absorption spectra which may be preferentially excited by our multispectral analysis. For example, absorption spectra for extracted zeaxanthin and violaxanthin pigments indicate that both absorb light from 420nm, 442nm, and 458nm excitation, but longer wavelength excitation (505nm and 525nm) may not be as readily absorbed by violaxanthin (Ruban et al., 2001). The spectrally-dependent variance in NPQ response observed for phenotypes 1 and 3 (Fig. 3i-k) may potentially reflect differences in the relative abundance and utilization of various xanthophyll pigments. While additional research is needed, such a connection between XC pigment pool/utilization and excitation wavelength could provide an additional dimension for understanding NPQ responses and how they might differ across species and environmental conditions.

Significant spectrally-dependent variability is also notable within the τ_1_ ^ST^ and τ_2_ ^ST^ reoxidation kinetics. These time constants reflect the rate of electron transport between the Qa and Qb site of the PSII reaction center (τ ^ST^) and further downstream kinetics involving the PQ pool (τ ^ST^) and further downstream electron transport. Previous work has indeed demonstrated the utility of τ ^ST^ for characterizing light or thermal acclimation state in reef corals (Suggett et al., 2022; Hoadley et al., 2019, Hoadley et al., 2023) or productivity rates in marine algae (Gorbunov & Falkowski, 2020). Values from τ ^ST^ are less well understood yet the clear structure observed in our profiles suggest this metric is indeed useful for assessing trait-based differences across species and/or environmental conditions.

### Linking primary cellular traits with photophysiology

Underlying *Symbiodiniaceae* cellular physiology differed significantly across the four phenotypes derived from chlorophyll-*a* fluorescence-based measurements. Linking underlying cellular physiology with more easily measured secondary traits such as photo physiology is critical for broadening the utility of multispectral and single-turnover chlorophyll-*a* fluorometers. These non-invasive, optical tools could serve as highly informative platforms for monitoring health and resilience of photosynthetic organisms, including reef corals (Suggett et al., 2022). Cellular traits, such as granularity which broadly measures the light scattering properties of a cell, were significantly higher in phenotypes 1 and 3 and may serve to deflect excess excitation energy. Reductions in photochemical quenching are more quickly relaxed in phenotypes 1 and 3 and higher granularity may serve to mitigate rapid shifts in light, functioning to reflect excess excitation energy away from the cell and reducing reliance on downstream processes such as closing PSII reaction centers in response to high light (qP, Fig. 3n, p). In contrast, large cell size and lower chlorophyll content cell^-1^ for phenotype 2 could reduce the package effect within these symbionts, thereby reducing overall reliance on NPQ (Fig. 3j) as less light is captured by each individual cell. Indeed, cellular characteristics may help explain the photo physiological strategies employed by each phenotype, and further correlative analysis between primary and secondary traits is warranted.

As chlorophyll-*a* fluorescence-based measurements are increasingly utilized for understanding photosynthetic poise and the utilization of stress response mechanisms by coral photosymbionts, identifying direct linkages between photo physiology and ecosystem services or underlying cellular traits are needed (Suggett et al., 2017). Our network analysis identified key correlations between basic cellular traits and photo physiological parameters across 20 different coral/symbiont combinations (Fig. 5). Certain chlorophyll-*a* fluorescence-derived photophysiological parameters may serve as useful biomarkers for some primary cellular traits, especially when properly contextualized within specific environmental and/or acclimatory conditions. For example, N:P ratios are inversely correlated to quantum yield of PSII measurements and directly correlated with τ ^ST^ reoxidation kinetics suggesting that phosphorous limited cells downregulate photochemical activity. Specifically, reductions in the quantum yield of PSII indicate reduced efficiency of light utilization for photochemistry whereas increases in τ ^ST^ reflect slower rates of electron transport, both of which appear to occur as N:P ratios rise. From a genotype perspective, both cell size and C:P differ significantly across the predominantly *Cladocopium C15* and predominantly *Durusdinium D1* phenotypes (phenotypes 1 and 2 respectively, Fig. 4a, e; Fig. 6a, c). These genotype level differences in cellular physiology may also be reflected in the reoxidation kinetics values where *Durusdinium D1* (phenotype 2) appears to have slower (higher rate constant) light acclimated τ ^ST^ (Fig. 3q, r) yet faster (lower rate constant) τ ^ST^ (Fig. 3u, v) reoxidation rates than those for *Cladocopium C15* (Fig. 6d). Linking both primary and secondary trait differences, especially across species known for their thermal tolerance, can be valuable for understanding what function traits are important for establishing resilient coral symbioses.

Carbon to phosphorous ratios also appear to be correlated with various photochemical metrics, most notably photochemical and non-photochemical quenching mechanisms which function to balance light utilization within the cell. Phosphorous limitation thus appears well linked to photochemical metrics, potentially regulating gene expression along with cell ultrastructure (Rosset et al., 2017; Ferrier-Pagès et al., 2016; Lin et al., 2019). Granularity is also linked with many different photochemical metrics which is perhaps not surprising given the strong phenotype differences observed for this cellular trait (Fig. 4). Overall, the strong linkages observed in our network analysis help strengthen our understanding of how differences in cellular traits across Symbiodiniaceae species regulate chlorophyll-*a* fluorescence-based phenotypes.

## Conclusions

The trends in this study further emphasize the utility of using photo-physiologically derived biomarkers across a variable light protocol to elicit different phenotypic responses in coral photosymbionts. Through the collection and analysis of large-scale chlorophyll fluorescence data sets, it is possible to resolve differences across *in hospite* coral symbionts for some species, regardless of growth environment. Further, by identifying correlations between critical first-order cellular traits and second-order photo physiological measurements, we can gain insight regarding how cellular mechanisms and characteristics affect algal photosynthesis under environmental stress. Implementation of low-cost, open-sourced methods of fluorescence measurement in coral restoration facilities may allow for quick determination of endosymbiont characteristics and better identification of the traits which underly thermal tolerance.

## Author Contributions

A.M and K.H. planned and designed the research. A.M., B.P., S.L., S.W., L.L., and K.H conducted fieldwork, and analyzed data. T.M maintained the coral in healthy and stable conditions prior to and during our experiment. A.M and K.H wrote the manuscript. All authors provided feedback on the manuscript. A.M. and K.D.H agree to serve as corresponding authors, responsible for contact and communication.

## Data Availability

All data needed to evaluate the conclusions in the paper are present in the paper and/or the Supplementary Materials. Pending scientific review, raw data and analytical scripts for Figs 2-5 will be available via github (khoadley/coral_phenotypes_2023).

## Supporting information

table for physiology

symportal data

