## Supplementary material for "Cellular traits regulate fluorescence-based bio-optical phenotypes of coral photosymbionts living *in-hospite*": table for physiology

**Supplemental Information**

**Supplemental Table 1: Average C, N, and P content and ratios for symbionts across coral hosts.** Each measurement is an average of 3 replicates, with +/- 1 standard deviation from the mean. Where a standard deviation is not present, fewer than 3 replicates were available.

|  | **Symbiont** | **Host** | **C per Cell**  **(pmol)** | **N per Cell**  **(pmol)** | **P per Cell**  **(pmol)** | **C:N** | **N:P** | **C:P** |
| --- | --- | --- | --- | --- | --- | --- | --- | --- |
| **Outdoor** | **C3** | *A.hum* var 1 | 136.37  +/- 20.72 | 20.57  +/- 4.99 | 1.26  +/- 0.31 | 6.71  +/- 0.56 | 16.49  +/- 0.81 | 110.59  +/- 8.85 |
|  | **C3** | *A.mill* | 322.46  +/- 159.63 | 41.80  +/- 23.22 | 1.15  +/- 0.66 | 7.87  +/- 0.96 | 27.34  +/- 1.31 | 228.66  +/- 29.17 |
|  | **C21** | *Acropora* sp | 151.57  +/- 96.62 | 21.13  +/- 13.84 | 0.90  +/- 0.05 | 7.29  +/- 0.47 | 22.58  +/- 14.30 | 161.77  +/- 98.5 |
|  | **C3** | *A.val* | 369.08  +/- 233.10 | 39.30  +/- 27.29 | 1.24  +/- 0.83 | 9.68  +/- 0.86 | 27.37  +/- 2.44 | 263.71 |
|  | **C21** | *A.yong* | 136.13  +/- 7.97 | 18.50  +/- 1.15 | 0.62  +/- 0.03 | 7.36  +/- 0.14 | 30.03  +/- 1.71 | 220.97  +/- 13.47 |
|  | **C1** | *C.chal* | 202.31  +/- 20.06 | 22.20  +/- 1.30 | 1.04  +/- 0.26 | 9.10  +/- 0.51 | 21.73  +/- 4.76 | 199.39  +/- 53.09 |
|  | **C26** | *M.cap* var 1 | 300.37  +/- 45.76 | 32.55  +/- 6.01 | 1.75  +/- 0.22 | 9.27  +/- 0.32 | 19.04  +/- 7.63 | 175.23  +/- 64.59 |
|  | **C15** | *M.cap* var 2 | 135.37  +/- 41.99 | 21.27  +/- 6.93 | 0.64  +/- 0.17 | 6.40  +/- 0.13 | 33.10  +/- 4.21 | 211.35  +/- 22.91 |
|  | **C1** | *M.cap* var 3 | 193.62  +/- 14.30 | 23.35  +/- 1.77 | 0.88  +/- 0.20 | 8.30 | 29.87  +/- 0.24 | 247.76  +/- 1.91 |
|  | **C15** | *M.dig* | 229.39  +/- 13.16 | 27.30  +/- 1.73 | 0.92 | 8.43  +/- 0.49 | 28.25  +/- 2.83 | 234.26  +/- 5.44 |
|  | **C1** | *P.cact* | 154.91  +/- 29.47 | 22.33  +/- 7.16 | 0.98  +/- 0.24 | 7.14  +/- 1.02 | 22.6  +/- 2.39 | 159.97  +/- 8.26 |
|  | **D1** | *P.cont* | 185.32  +/- 4.30 | 26.93  +/- 4.71 | 1.16  +/- 0.13 | 7.00  +/- 1.02 | 25.10  +/- 1.78 | 171.41  +/- 23.12 |
|  | **D1** | *P.dam* var 1 | 204.08  +/- 104.56 | 28.83  +/- 13.49 | 2.04 | 6.99  +/- 0.33 | 21.54 | 157.51 |
|  | **D1** | *P.dam* var 2 | 257.14  +/- 42.22 | 37.10  +/- 7.23 | 1.82  +/- 0.34 | 6.96  +/- 0.25 | 20.27  +/- 1.01 | 141.02  +/- 6.67 |
|  | **D1** | *T.ren* var 1 | 168.46 | 28.70 | 1.12 | 5.52  +/- 0.98 | 29.73  +/- 5.28 | 161.05  +/- 14.54 |
|  | **D1** | *T.ren* var 2 | 414.87  +/- 44.20 | 60.00  +/- 9.31 | 2.32  +/- 0.54 | 6.95  +/- 0.44 | 24.79  +/- 1.25 | 177.91  +/- 16.38 |
| **Indoor** | **C3** | *A.hum* var 2 | 182.29  +/- 39.75 | 28.63  +/- 1.22 | 1.51  +/- 0.38 | 6.34  +/- 1.17 | 20.17  +/- 4.02 | 124.93  +/- 6.74 |
|  | **C15** | *M.cap* var 2 | 171.06  +/- 37.21 | 22.83  +/- 3.78 | 0.70  +/- 0.08 | 7.45  +/- 0.40 | 30.58  +/- 3.55 | 221.02  +/- 18.15 |
|  | **C15** | *M.cap* var 3 | 171.80  +/- 23.94 | 18.75  +/- 2.05 | 0.65  +/- 0.12 | 9.31  +/- 2.34 | 28.78  +/- 6.28 | 260.64  +/- 9.00 |
|  | **D1** | *T.ren* var 2 | 113.50  +/- 59.54 | 15.47  +/- 8.17 | 0.75  +/- 0.48 | 7.34  +/- 0.04 | 23.96  +/- 8.26 | 175.95  +/- 60.91 |

**Supplemental Table 2: Average values for cellular metrics determined via flow cytometry for *Symbiodiniaceae* across coral hosts.** Each measurement is an average of 3 replicates, with +/- 1 standard deviation from the mean. Where a standard deviation is not present, fewer than 3 replicates were available for that metric. Neutral lipid content is reported in Fluorescence Units (FU).

|  | **Symbiont** | **Host** | **Neutral Lipids**  **(FU)** | **Bead-Norm**  **Chl *a*** | **Bead-Norm**  **FSC (Cell Size)** | **Bead-Norm**  **SSC (Granularity)** |
| --- | --- | --- | --- | --- | --- | --- |
| **Outdoor** | **C3** | *A.hum* var 1 | 1418.67  +/- 716.38 | 6.32  +/- 0.11 | 3.67  +/- 0.28 | 1.96  +/- 0.14 |
|  | **C3** | *A.mill* | 477.33  +/- 826.77 | 20.39  +/- 0.17 | 3.32  +/- 0.17 | 2.55  +/- 0.09 |
|  | **C21** | *Acropora* sp | 3015.00  +/- 1727.71 | 26.83  +/- 1.85 | 3.60  +/- 0.40 | 1.49  +/- 0.03 |
|  | **C3** | *A.val* | 1894.00  +/- 934.05 | 21.35  +/- 1.26 | 3.61  +/- 0.16 | 3.08  +/- 0.41 |
|  | **C21** | *A.yong* | 6606.00  +/- 7338.48 | 25.95  +/- 1.20 | 3.77  +/- 0.06 | 2.01  +/- 0.09 |
|  | **C1** | *C.chal* | 6047.67  +/- 540.51 | 20.52  +/- 0.58 | 3.73  +/- 0.15 | 2.53  +/- 0.05 |
|  | **C26** | *M.cap* var 1 | 2461.67  +/- 1118.45 | 30.65  +/- 0.65 | 3.74  +/- 0.59 | 2.86  +/- 0.43 |
|  | **C15** | *M.cap* var 2 | 2352.33  +/- 668.13 | 87.33  +/- 110.69 | 3.78  +/- 0.10 | 3.16  +/- 0.39 |
|  | **C1** | *M.cap* var 3 | 1895.33  +/- 710.88 | 23.92  +/- 1.39 | 3.79  +/- 0.12 | 3.39  +/- 0.31 |
|  | **C15** | *M.dig* | 1749.00  +/- 248.57 | 21.60  +/- 0.39 | 3.66  +/- 0.12 | 2.84  +/- 0.16 |
|  | **C1** | *P.cact* | 2049.33  +/- 1776.15 | 20.11  +/- 0.55 | 3.89  +/- 0.19 | 2.02  +/- 0.11 |
|  | **D1** | *P.cont* | NA | 21.47  +/- 0.14 | 4.52  +/- 0.42 | 2.91  +/- 0.30 |
|  | **D1** | *P.dam* var 1 | 7640.33  +/- 1245.23 | 18.19  +/- 0.85 | 4.22  +/- 0.59 | 2.25  +/- 0.67 |
|  | **D1** | *P.dam* var 2 | 7389.33  +/- 6651.45 | 22.10  +/- 2.02 | 4.68  +/- 0.80 | 2.42  +/- 0.30 |
|  | **D1** | *T.ren* var 1 | 10020.00 | 20.06 | 5.21 | 3.79 |
|  | **D1** | *T.ren* var 2 | 1652.00  +/- 2861.35 | 22.68  +/- 1.53 | 5.34  +/- 0.51 | 3.75  +/- 0.30 |
| **Indoor** | **C3** | *A.hum* var 2 | 955.67  +/- 460.65 | 21.44  +/- 0.87 | 3.23  +/- 0.09 | 2.57  +/- 0.23 |
|  | **C15** | *M.cap* var 2 | 2248.33  +/- 1143.01 | 19.93  +/- 0.95 | 3.67  +/- 0.12 | 2.69  +/- 0.13 |
|  | **C15** | *M.cap* var 3 | 1485.00  +/- 81.07 | 22.41  +/- 0.27 | 3.49  +/- 0.07 | 3.68  +/- 0.03 |
|  | **D1** | *T.ren* var 2 | 7916.67 | 18.55  +/- 0.18 | 4.20  +/- 0.03 | 2.54  +/- 0.01 |

**Supplemental Table 3: Bonferroni-adjusted p-values comparing photophysiology across excitation wavelength within each of the 4 phenotypic profiles.** These statistics correspond with the phenotypic profiles found in Figure 3. Significant comparisons across excitation wavelengths appear in bold.

|  | λ (nm) | | λ (nm) | Φ_PSII_ | σ_PSII_ | qP | NPQ | τ_1_ | τ_2_ |
| --- | --- | --- | --- | --- | --- | --- | --- | --- | --- |
| Phenotype 1 | 442 | | 420 | 4.05E-01 | **<2e-16** | 1.000 | **2.69E-12** | **<2e-16** | 1 |
|  | 458 | | 420 | **2.22E-15** | **<2e-16** | 1.000 | **2.00E-16** | **<2e-16** | **2.00E-16** |
|  | 505 | | 420 | 1 | **<2e-16** | 1.000 | **2.00E-16** | **<2e-16** | **2.00E-16** |
|  | 525 | | 420 | **2.22E-15** | **<2e-16** | 1.000 | **2.00E-16** | **<2e-16** | **2.00E-16** |
|  | 458 | | 442 | **2.00E-16** | **<2e-16** | 1.000 | **0.000126** | **<2e-16** | **2.00E-16** |
|  | 505 | | 442 | **0.00432** | **<2e-16** | 1.000 | **1.33E-11** | **<2e-16** | **2.00E-16** |
|  | 525 | | 442 | **2.00E-16** | **<2e-16** | 1.000 | **6.09E-05** | **<2e-16** | **2.00E-16** |
|  | 505 | | 458 | **1.97E-10** | **<2e-16** | 1.000 | 6.45E-02 | **<2e-16** | **4.60E-08** |
|  | 525 | | 458 | 1 | **<2e-16** | 1.000 | 1 | **<2e-16** | 1 |
|  | 525 | | 505 | **1.76E-10** | **<2e-16** | 1.000 | 1.03E-01 | 1 | **8.44E-11** |
| Phenotype 2 | 442 | | 420 | **5.60E-03** | **2.00E-16** | 1.000 | **1.45E-12** | **2.00E-16** | 1 |
|  | 458 | | 420 | 1 | **2.00E-16** | 1.000 | **2.00E-16** | **2.00E-16** | **2.00E-16** |
|  | 505 | | 420 | **1.24E-06** | **2.00E-16** | 2.66E-01 | **2.00E-16** | **2.00E-16** | **0.015262** |
|  | 525 | | 420 | 1 | **2.00E-16** | 1.000 | **0.007534** | **2.00E-16** | **2.00E-16** |
|  | 458 | | 442 | **1.33E-03** | **2.00E-16** | 1.000 | 1 | **2.00E-16** | **2.00E-16** |
|  | 505 | | 442 | 0.6621 | **2.00E-16** | 1.000 | **0.001224** | **2.00E-16** | **2.16E-05** |
|  | 525 | | 442 | **1.19E-02** | **2.00E-16** | 1.000 | **0.000576** | **2.00E-16** | **2.00E-16** |
|  | 505 | | 458 | **1.53E-07** | **4.64E-08** | 1.000 | 0.056438 | **0.00616** | **2.00E-16** |
|  | 525 | | 458 | 1 | **2.00E-16** | 1.000 | **3.47E-06** | **4.44E-15** | **0.000259** |
|  | 525 | | 505 | **3.80E-06** | **2.00E-16** | 1.000 | **3.77E-14** | **2.88E-05** | **2.00E-16** |
| Phenotype 3 | 442 | | 420 | **7.59E-06** | **2.00E-16** | 1 | **2.00E-16** | **2.88E-07** | **0.00121** |
|  | 458 | | 420 | **0.00698** | **2.00E-16** | 1 | **2.00E-16** | **2.00E-16** | 1 |
|  | 505 | | 420 | **2.00E-16** | **2.00E-16** | 0.799 | **2.00E-16** | **2.00E-16** | **1.73E-07** |
|  | 525 | | 420 | **1.21E-06** | **2.00E-16** | 1 | **2.00E-16** | **2.00E-16** | 1 |
|  | 458 | | 442 | 1 | **2.00E-16** | 1 | 0.3355 | **1.13E-13** | **0.00421** |
|  | 505 | | 442 | **0.00754** | **2.00E-16** | 0.93 | **4.34E-06** | **2.00E-16** | 0.73067 |
|  | 525 | | 442 | 1 | **2.00E-16** | 1 | 1 | **2.00E-16** | **0.00227** |
|  | 505 | | 458 | **8.46E-06** | **0.000476** | 1 | **0.0341** | **0.02048** | **1.04E-06** |
|  | 525 | | 458 | 0.57103 | **2.00E-16** | 1 | 0.4726 | **8.59E-10** | 1 |
|  | 525 | | 505 | **0.02512** | **2.00E-16** | 1 | **9.02E-06** | **0.00657** | **4.26E-07** |
| Phenotype 4 | 442 | | 420 | **2.90E-02** | **<2e-16** | 1 | **3.69E-04** | **2.00E-16** | 1 |
|  | 458 | | 420 | 1 | **<2e-16** | 1 | **5.78E-08** | **2.00E-16** | **<2e-16** |
|  | 505 | | 420 | **1.61E-07** | **<2e-16** | 1.62E-01 | **2.00E-16** | **2.00E-16** | 1 |
|  | 525 | | 420 | 1 | **<2e-16** | 5.48E-01 | **1.73E-05** | **2.00E-16** | **<2e-16** |
|  | 458 | | 442 | **3.32E-03** | **<2e-16** | 1 | 0.897146 | **2.00E-16** | **<2e-16** |
|  | 505 | | 442 | 0.075415 | **<2e-16** | 1 | **3.99E-09** | **2.00E-16** | 5.02E-01 |
|  | 525 | | 442 | 1 | **<2e-16** | 1 | 1 | **2.00E-16** | **<2e-16** |
|  | 505 | | 458 | **3.82E-09** | **2.09E-02** | 1 | **5.17E-05** | **1.83E-08** | **<2e-16** |
|  | 525 | | 458 | 4.40E-01 | **<2e-16** | 1 | 1 | **2.00E-16** | 1 |
|  | 525 | | 505 | **2.16E-04** | **<2e-16** | 1 | **2.16E-07** | 1.40E-01 | **<2e-16** |

**Supplemental Table 4: Bonferroni-adjusted p-values comparing photophysiology across phenotype for each excitation wavelength.** These statistics correspond with the phenotypic profiles found in Figure 3. Significant comparisons appear in bold.

|  | Phen | | Phen | Φ_PSII_ | σ_PSII_ | qP | NPQ | τ_1_ | τ_2_ |
| --- | --- | --- | --- | --- | --- | --- | --- | --- | --- |
| 420 nm | 2 | | 1 | **1.21E-09** | 1 | **1.30E-07** | **2.00E-16** | **1.13E-03** | **2.30E-10** |
|  | 3 | | 1 | 1 | 1 | 1 | **6.97E-06** | 4.58E-01 | 1 |
|  | 4 | | 1 | **6.05E-12** | 1 | **1.49E-05** | **3.59E-11** | 5.53E-02 | **7.43E-09** |
|  | 3 | | 2 | **2.93E-05** | 0.353 | **0.000226** | **2.33E-05** | 1 | **2.24E-04** |
|  | 4 | | 2 | 1 | 1 | 1 | **5.04E-07** | 1 | 1 |
|  | 4 | | 3 | **2.37E-06** | 1 | **0.005604** | 1 | 1 | **0.002287** |
| 442 nm | 2 | | 1 | **2.66E-15** | 1 | **1.99E-07** | **2.00E-16** | **3.87E-04** | **4.39E-12** |
|  | 3 | | 1 | **0.015948** | 1 | 1 | **3.70E-09** | **1.04E-02** | **2.15E-03** |
|  | 4 | | 1 | **2.00E-16** | 1 | **1.19E-05** | **6.38E-09** | 1.32E-01 | **8.86E-11** |
|  | 3 | | 2 | **0.001686** | 1 | **0.00302** | **0.00126** | 1 | 0.13268 |
|  | 4 | | 2 | 1 | 1 | 1 | **1.52E-10** | 3.28E-01 | 1 |
|  | 4 | | 3 | **0.000253** | 1 | **0.03625** | 0.78251 | 0.956649 | 0.51449 |
| 458 nm | 2 | | 1 | **2.00E-16** | 1 | **2.44E-08** | **2.00E-16** | 1.41E-01 | **1.55E-12** |
|  | 3 | | 1 | **4.38E-06** | 1 | 1 | **3.93E-07** | **1.28E-04** | **1.60E-06** |
|  | 4 | | 1 | **2.00E-16** | 0.433 | **1.02E-05** | **1.83E-05** | 8.81E-02 | **2.43E-12** |
|  | 3 | | 2 | 0.0963 | 1 | **8.46E-05** | **4.84E-03** | 9.79E-02 | 1 |
|  | 4 | | 2 | 1 | 1 | 0.94526 | **1.98E-10** | 1 | 1 |
|  | 4 | | 3 | **0.0435** | 1 | **0.00465** | 0.38599 | 0.087283 | 1 |
| 505 nm | 2 | | 1 | **2.00E-16** | 1 | **2.27E-08** | **2.00E-16** | **3.06E-03** | **3.21E-11** |
|  | 3 | | 1 | **2.54E-06** | 1 | 1 | **9.47E-07** | **3.16E-06** | **1.11E-06** |
|  | 4 | | 1 | **2.00E-16** | 1 | **1.45E-05** | **2.16E-05** | **2.03E-02** | **9.93E-12** |
|  | 3 | | 2 | 0.3312 | 1 | **2.21E-05** | **2.80E-02** | 1.76E-01 | 1 |
|  | 4 | | 2 | 1 | 1 | 0.80667 | **4.28E-08** | 1 | 1 |
|  | 4 | | 3 | **0.0407** | 0.491 | **0.00198** | 0.521 | **0.02679** | 1 |
| 525 nm | 2 | | 1 | **2.00E-16** | 1 | **1.47E-08** | **2.00E-16** | 1.20E-01 | **1.73E-14** |
|  | 3 | | 1 | **1.78E-07** | 1 | 1 | **2.43E-04** | **1.47E-05** | **3.05E-08** |
|  | 4 | | 1 | **2.00E-16** | 1 | **2.05E-05** | **1.22E-03** | **4.40E-02** | **1.33E-15** |
|  | 3 | | 2 | 0.7822 | 1 | **7.93E-06** | **2.82E-02** | **2.94E-02** | 1 |
|  | 4 | | 2 | 1 | 1 | 0.60249 | **9.95E-07** | 1 | 1 |
|  | 4 | | 3 | 0.0979 | 0.115 | **0.00135** | 1 | **0.0383** | 1 |
